# Acute ampakines increase voiding function and coordination in a rat model of SCI

**DOI:** 10.1101/2023.05.26.542339

**Authors:** Sabhya Rana, Firoj Alom, Robert C Martinez, David D Fuller, Aaron D Mickle

## Abstract

Neurogenic bladder dysfunction causes urological complications and reduces the quality of life in persons with spinal cord injury (SCI). Glutamatergic signaling via AMPA receptors is fundamentally important to the neural circuits controlling bladder voiding. Ampakines are positive allosteric modulators of AMPA receptors that can enhance the function of glutamatergic neural circuits after SCI. We hypothesized that ampakines can acutely stimulate bladder voiding that has been impaired due to thoracic contusion SCI. Adult female Sprague Dawley rats received a unilateral contusion of the T9 spinal cord (n=10). Bladder function (cystometry) and coordination with the external urethral sphincter (EUS) were assessed five days post-SCI under urethane anesthesia. Data were compared to responses in spinal intact rats (n=8). The “low impact” ampakine CX1739 (5, 10, or 15 mg/kg) or vehicle (HPCD) was administered intravenously. The HPCD vehicle had no discernable impact on voiding. In contrast, following CX1739, the pressure threshold for inducing bladder contraction, voided volume, and the interval between bladder contractions were significantly reduced. These responses occurred in a dose-dependent manner. We conclude that modulating AMPA receptor function using ampakines can rapidly improve bladder voiding capability at sub-acute time points following contusion SCI. These results may provide a new and translatable method for therapeutic targeting of bladder dysfunction acutely after SCI.

## Introduction

Traumatic spinal cord injury (SCI) often results in bladder dysfunction which can lead to bladder infection and reduced quality of life (Hamid et al., 2018; Piatt et al., 2016). Indeed, restoration of bladder function is ranked as one of the highest priorities by individuals with SCI (Bourbeau et al., 2020; French et al., 2010). The fundamental problem is uncoordinated voiding due to damage to sensory and motor circuits that control bladder coordination. The symptoms can include urinary retention (hypoactive) or incontinence (hyperactive) depending on the injury and time after recovery, leading to urodynamic complications such as urinary infection, kidney damage, bladder cancer, urethral strictures, and bladder stones (Taweel and Seyam, 2015).

The synaptic pathways which serve to coordinate the activation of bladder detrusor and external urethral sphincter (EUS) muscles rely on glutamatergic neurotransmission, and in SCI, these circuits can be disrupted (de Groat et al., 2015; Fowler et al., 2008; Yoshiyama, 2009; Yoshiyama et al., 1999b). The α-amino-3-hydroxy-5-methyl-4-isoxazolepropionic acid (AMPA) glutamate receptor plays a significant role in mediating drive in the intact micturition reflex (Yoshiyama and de Groat, 2005; Yoshiyama et al., 1997). After SCI, the bladder is initially in an areflexive state, and as spinal circuits are reorganized, the bladder transitions to neurogenic detrusor overactivity (de Groat et al., 2015; de Groat and Yoshimura, 2006). Glutamatergic signaling and AMPA receptors are altered during these states (Grossman et al., 2001; Grossman et al., 1999; Mitsui et al., 2011; Pikov and Wrathall, 2001, 2002). In general, there are decreases in AMPA receptor subunits initially after an injury during the hypoactive bladder state (Grossman et al., 1999).

The AMPA receptor can be allosterically modulated using pharmacologic compounds called ampakines (Arai and Kessler, 2007), and recent work suggests that these compounds have therapeutic value after SCI (Rana et al., 2021; Wollman et al., 2020). For example, when given after acute or chronic SCI in rats, ampakines have a powerful, beneficial impact on the glutamatergic synaptic pathways, which control breathing (Rana et al., 2021; Wollman et al., 2020). After high cervical SCI, diaphragm output is diminished, but systemically providing a low dose of ampakine can cause a rapid and sustained increase in diaphragm electromyogram output (Rana et al., 2021). Since depolarization of the phrenic motoneurons that innervate the diaphragm critically depends on AMPA receptor activation (Chitravanshi and Sapru, 1996; Fuller et al., 2022; Rana et al., 2019), ampakines are likely acting at least in part on spinal circuits to stimulate diaphragm output (Thakre et al., 2022).

Building on the foundation of prior work in ampakine treatment for respiratory insufficiency, we studied the impact of ampakines on voiding reflexes after SCI. An adult rat model of sub-acute (5-days) thoracic contusion SCI was used to test the hypothesis that intravenous delivery of ampakine CX1739 would have a dose-dependent ability to restore bladder and EUS functions and their coordinated voiding. Bladder function was evaluated by continuous flow cystometry (Andersson et al., 2011). The results demonstrate a remarkable ability of ampakine treatment to restore voiding reflexes following SCI. Ampakines’ rapid and powerful impact on bladder function provides a foundation for a new and translatable method for therapeutic targeting of bladder dysfunction after SCI.

## Methods

### Experimental Animals

A total of 18 adult (12 weeks old) female Sprague-Dawley rats (Hsd:Sprague Dawley® SD, Envigo Indianapolis, IN) were used in these studies. Most previous rat studies on SCI and bladder function have used females since there are considerably fewer post-operative complications and due to the ease of post-operative manual voiding when the rats are unable to self-void (Lin et al., 2016; Mitsui et al., 2014; Pikov and Wrathall, 2001). Our initial preliminary experiments included male and female rats however, only female rats recovered cystometric voiding 5 days after injury. For these reasons, female rats were chosen for this study.

All procedures were approved by the Institutional Animal Care and Use Committee at the University of Florida and are in accordance with the National Institutes of Health Guidelines. Animals were housed individually in cages under a 12-hr light/dark cycle with *ad libitum* access to food and water.

### T9 Contusion Injury

A cohort of rats in this study received a midline T9 contusion injury (n=10). Depth of ketamine (90 mg/kg) and xylazine (10 mg/kg) anesthesia was confirmed at the onset of any procedure using a lack of hind limb withdrawal or whisker twitch following a toe pinch. An adequate level of anesthesia was continuously verified during the surgical procedure, and the respiratory rate was measured every 15 mins. The animal was re-dosed with a 1/3 dose of the initial ketamine/xylazine dose if needed. Body temperature was continuously monitored using a rectal temperature probe and maintained between 37-38°C using a water-recirculating heating pad. Under sterile conditions, a dorsal incision was made from the T5-T11 region of the spine. Dorsal paravertebral muscles between T6-T10 were incised and retracted. The posterior portion of thoracic vertebrae was exposed T8-T10, and a laminectomy was performed at T9 while preserving the facet joints and leaving the dura intact. Rats were suspended by clamps secured laterally at the T8 and T10 vertebrae. A 2.5 mm impactor tip was aligned at the midline. Subsequently, rats were subjected to a single contusion injury. A desired force of 100 kDy with 0 s dwell time was delivered using the Infinite Horizon Impactor (Precision Systems and Instrumentation, Lexington, KY). Animals were de-clamped and the overlying muscles were sutured with sterile 4–0 webcryl. The overlying skin was closed using 9 mm wound clips. Animals were maintained on a heating pad until alert and awake. Animals received one dose of the extended-release buprenorphine SR LAB (1 mg/kg, ZooPharm) followed by carprofen (5 mg/kg, q.d.), baytril (5 mg/kg, q.d.), and lactated Ringer’s solution (10 ml/day, q.d.) for the initial 48 hr post-injury. Animals were monitored daily for signs of distress, dehydration, and weight loss, with appropriate veterinary care given as needed. No animals needed to be excluded from the study based on pre-determined exclusionary factors of bone-hit or slip tip during contusion, >10% difference in actual delivered force, limb autophagia or weight loss >20% of a pre-injury time point.

### In Vivo Bladder Cystometry Recordings

Five days following spinal cord injury or in intact rats, a 24 G *i.v.* catheter (SR#FF2419, Terumo Corporation) was placed on the distal end of the tail vein on the day of recording under 2% isoflurane. Rats were slowly infused with urethane (0.84 g/kg; 102452447, Sigma) (diluted in saline at 150 mg/mL) through the tail vein. Later 0.12 to 0.24 g/kg of urethane was given if required during cystometry, depending on animal’s response. An injection of saline was given subcutaneously (1 mL/100 gm of rat) to keep the rat hydrated during the recording time of cystometry under urethane. The abdominal area and the area around the base of the tail were shaved. The rat was then transferred to a closed loop heating pad to control the body temperature at 37°C (2221962P, CWE, Inc.). The rat was placed under 1-2% isoflurane (J121008, Akron, Inc.) through a nose cone. An incision was made in the abdominal wall, and a purse-string suture was made using a 5-0 nylon suture (07-809-8813, Patterson Veterinary) around the apex of the bladder. The apex of the bladder inside the purse string suture was cut, and a flared catheter (BB31695-PE/3, Scientific Commodities, Inc.) was inserted into the bladder lumen. The purse string suture was secured tightly around the catheter, and the bladder was checked for leaks with a brief saline infusion. The polyethylene catheter was connected with a syringe pump (GT1976 Genie Touch, Kent Scientific Corporation) to infuse the saline into the bladder at 6 mL per hour. The intraluminal pressure measurements were acquired by using an inline pressure transducer (503067, World Precision Instruments) connected to a Transbridge amplifier (TBM4-D, World Precision Instruments) and processed at a 1000 Hz sampling rate using the Micro 1401 (Cambridge Electronic Design) data acquisition system and Spike2 Software (V10; Cambridge Electronic Design). The muscle of the abdominal cavity and skin incisions were sutured with nylon suture, and the isoflurane was then lowered to zero.

Next, two Teflon insulated silver wires (570742, A-M Systems) were placed into the EUS muscle (5 mm of bare wire exposed) to measure sphincter EMG. Two 25 G needles containing the EMG wires were inserted into the skin and muscle at a distance of 3-5 mm from the urethral opening, and the other end of the wire was connected with the pre-amplifier (HZP, Grass, Astro-Med. Inc.) and then amplifier (RPS107E, Grass, Astro-Med Inc.). A third wire was inserted into the skin of the abdominal area of the body by using a 18 G needle and connected to the pre-amplifier as a ground. The EUS EMG data from the amplifier was processed at 20,000 Hz using the Micro 1401 data acquisition system and Spike 2 software (**Figure 1A**). 30 minutes after stopping isoflurane we started the saline infusion and cystometric pressure recording. The voids were collected in a weigh boat by placing the weigh boat under the urethral opening to collect the voided volume. These volumes were weighed on a balance and recorded after each cystometric void.

**Figure 1.**
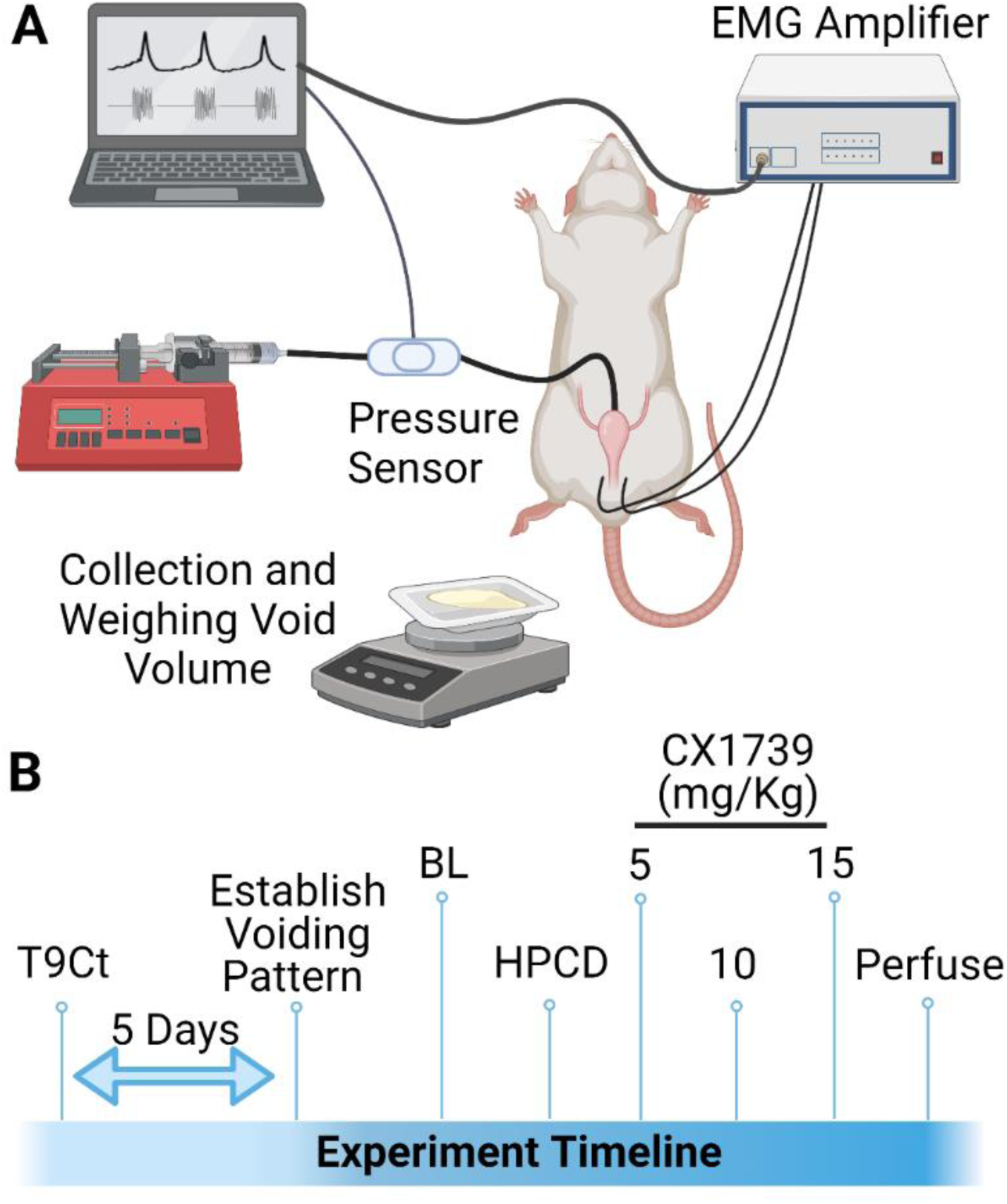
Experimental Setup and Design. A) Schematic of experimental urodynamic recordings in anesthetized rats. Volume of voided urine was also collected. B) Experimental timeline of studies. Animals received a unilateral T9 contusion, and urodynamic recordings were conducted at 5 days post contusion. During the recording, a baseline was established for urodynamic output. Animals received vehicle HPCD, and consecutive doses of ampakines CX1739 at doses of 5 mg/kg, 10 mg/kg, and 15 mg/kg. Each drug dose was spaced apart by 45 mins.

### Chemical and Pharmacological Agents

Ampakine CX1739 was provided by RespireRX. The drug was diluted in HPCD (Cat # H107-100G, Sigma) solution (10%) at a soluble concentration of 5 mg/mL. Aliquots were stored at −20°C for up to 6 months. Aliquots were thawed to room temperature on the day of each experiment. CX1739 was infused at concentrations of 5, 10, and 15 mg/kg intravenously slowly (∼2 mins). We then assessed the effects of those doses of CX1739 over the next 45 mins. Cystometry parameters were recorded as baseline, vehicle, 5 mg/kg, 10 mg/kg, and 15 mg/kg CX1739 in the same order (**Figure 1B**). Pharmacokinetic parameters for CX1739 in Sprague Dawley rats have previously been determined following an intravenous administration of CX1739. The mean plasma half-life of CX1739 was 1.25 ± 0.03 hrs, with a T_max_ of 30 minutes (information provided through personal communication with RespireRx). Although the 45 minutes interval between doses would not be within the time frame of *complete* post-administration clearance of the first CX1739 dose from the system, the plasma levels would be predicted to be considerably lower by 45 mins post administration. A limitation of terminal cystometry preparations is that the duration which an experiment in an anesthetized rat can be sustained is limited. In our experience recordings beyond six hours will dramatically increase variability in the data. The 45 min window allowed for the procedure to remain under ∼6 hours. Further, in our published studies investigating the impact of ampakines on breathing in rats following an SCI, the acute impact of intravenous ampakine administration wanes after approximately 30 minutes (Rana et al., 2021) Thus, the dosing schedule was based on three factors: 1) the drug ½ life; 2) the duration of which the experimental preparation was viable, and 3) data from the respiratory response of ampakines after SCI.

### Immunohistochemistry and Imaging

Following the cystometry experiments, animals were injected intraperitoneally with ketamine/xylazine cocktail (0.05 ml/100 gm) and transcardially perfused with 4% paraformaldehyde in 0.1 M phosphate-buffered saline (pH 7.4). T8-T10 spinal segments were resected, post-fixed for 24 hrs in 4% paraformaldehyde and cryopreserved in 30% sucrose in 0.1 M PBS (pH 7.4) for 3 days at 4°C. A 5 mm spinal cord segment centered at the injury epicenter was embedded in OCT and subsequently sectioned at 20 μm thickness in the transverse plane. Rehydrated sections were stained with 0.1% cresyl violet in glacial acetic acid. After 3 minutes, slides were rinsed in water and dehydrated through graded alcohol steps followed by xylene. Tissue sections were imaged and stitched using a 10x objective on a Keyence microscope (BZ-X700, Keyence Corporation of America, Itasca, IL).

### Data Analysis

Animals that did not develop a regular voiding pattern were not included in the group analysis as cystometric properties could not be measured. 3 out of 10 SCI rats did not develop a normal pattern, all intact rats exhibited a regular pattern (8 out of 8). These data were analyzed separately making comparisons within animals (**Figure 6**). Cystometric data (intercontraction interval, threshold pressure, peak pressure and baseline pressure as defined by (Andersson et al., 2011) was first analyzed by cystometry analyzer software (https://github.com/Samineni-Lab/cystometry_analyzer). Threshold pressures were verified manually to ensure accuracy. The cystometric voided saline was collected in a weigh boat and weighed on a balance to determine the volume of the voids. The total calculated volume of voids over a period of 30 min after the application of HPCD/CX1739 was divided by the total number of voids to determine the average voided volume. EUS EMG data was analyzed on Spike2 software (CED). The EMG signal was rectified, moving-median filtered using a 50 ms time constant, and finally smoothed using a 50 ms time constant to obtain the RMS EMG signal. The peak RMS EMG signal averaged over the duration of the burst to obtain RMS_peak_ EMG. The threshold activation pressure for EUS EMG was defined as the bladder pressure at which EUS EMG burst was evoked (intersection of red dotted lines in Figure 5A).

### Statistical Analysis

All statistical evaluations were performed using JMP statistical software (version 14.0, SAS Institute Inc., Cary, NC). Power calculations for each cohort of animals in the study were determined before the initiation of the study and based on pilot experiments not included in this article. The study statistical design (n=8 rats/group) was powered to consider the expected standard deviation in cystometrogram threshold pressure at 5 days post-contusion (22%) and in order to detect a 25% difference at a power of 0.8 and an α of 0.05. Using the mixed linear model, statistical significance was established at the 0.05 level and adjusted for any violation of the assumption of sphericity in repeated measures using the Greenhouse–Geisser correction. Group (Intact or Injured) and treatment (baseline, HPCD, CX1739 5 mg/kg, CX1739 10 mg/kg, and CX1739 15 mg/kg) were included as model variables, using animals as a random effect. When appropriate, *post-hoc* analyses were conducted using Tukey–Kramer honestly significant difference (HSD). Normality of the distribution was assessed using the Shapiro–Wilk test within each animal. Outliers were identified for each animal using an outlier box plot. No data were excluded from the analysis to highlight the responders versus the non-responders in each treatment group. Each data point on the graphs represents as mean ± SEM. Figures were designed and produced by using JMP and Adobe Illustrator.

## Results

### T9 Contusion Injury

A T9 midline contusion injury was effectively delivered in ten rats, with a distinct hematoma visually evident under the surgical microscope. The average impact force for the contusion group was 104 ± 4 kDy, and the average displacement was 582 ± 102 μm. All animals recovered successfully from the surgery. Following surgery, all animals in the contusion group exhibited bilateral hind-limb motor deficits. At the terminal experiment, upon dissection of the spinal cord, the level of injury was also confirmed in all animals by locating the T9 vertebral level and dorsal root relationship to the bruise. Five days post contusion, animals lost ∼5% of initial body weight (241 ± 20 pre-injury vs 228 ± 17 at 5 days post injury).

Representative histological images of the T9 spinal segment from an intact and contused animal are provided in **Figure 2**. Cross sections from the injury epicenter in injured animals demonstrated extensive pathology of the dorsal to ventral grey matter regions and dorsal white matter tracts. There was some residual sparing of lateral and ventral white matter tracts at the injury epicenter.

**Figure. 2.**
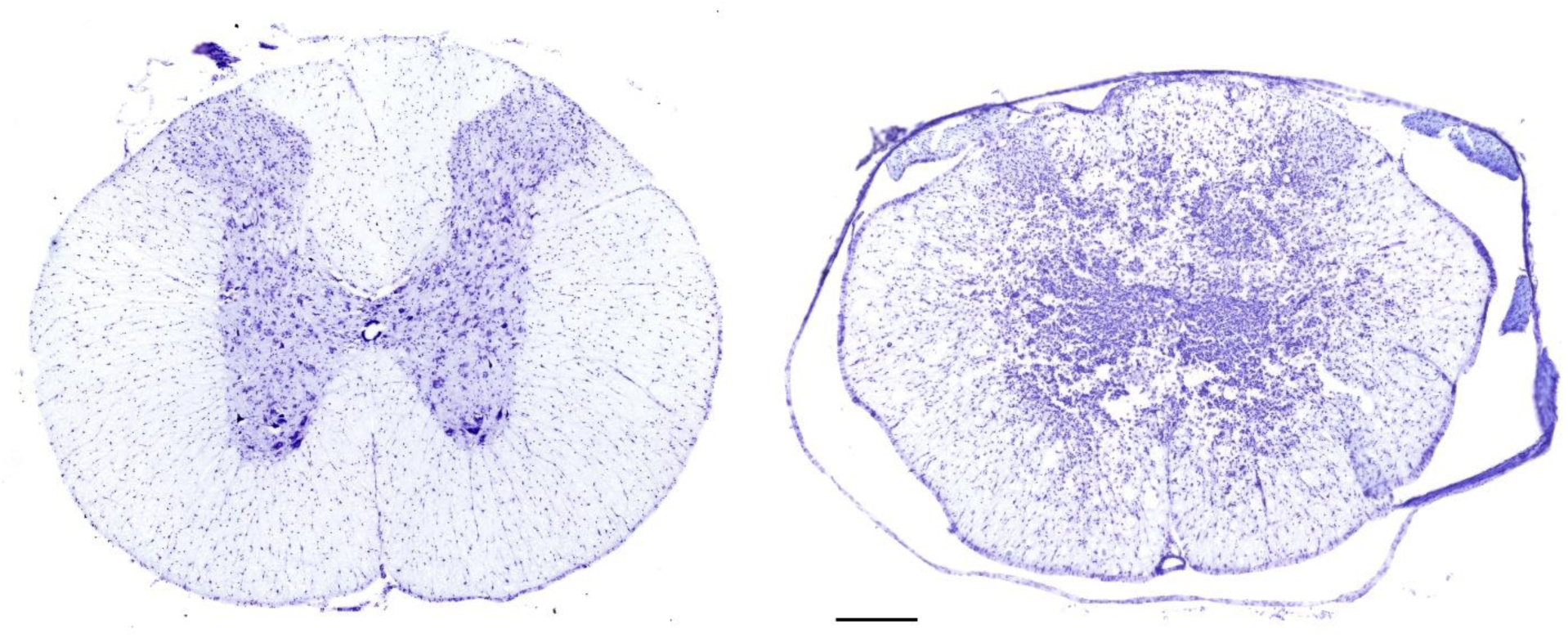
Histological assessment of T9 Contusion (T9Ct) injury epicenter. Serial, transverse sections (20 μm thickness) spanning ∼1 cm around the lesion epicenter (T9-T10) were stained with Cresyl violet in order to identify the extent of injury. A representative image from an intact (left) and T9 contused (right) spinal cord. Scale bar, 300 μm.

### Effect of Injury on Cystometric Bladder Function and EUS EMG Activity

Following the T9 contusion injury, rats acutely lost the ability to micturate spontaneously. Bladders were manually expressed 2-3 times daily for the first 3 days post-injury. Urodynamic recordings were performed 5 days post contusion in injured rats and compared to spinal-intact rats. Representative urodynamic recordings are presented in **Figure 3A**. Coordinated bladder contractions and associated EUS EMG activity were readily demonstrated in all 8 naïve animals. Cystometrogram recordings were characterized by a low baseline bladder volume during the saline-filling phase. As saline infusion reached the threshold volume, a bladder contraction occurred that was demonstrated by a pronounced rapid and brief increase in intravesical pressure and subsequently followed by voiding. Tonic EUS EMG activity was observed before the onset of voiding, followed by coordinated EUS bursting activity during the voiding phase. Distinct active and silent periods could be identified during the coordinated EUS “bursting” phase, which coincided with intravesical pressure oscillations in the cystometrogram. By this time point, 7 out of 10 injured rats exhibited impaired but spontaneous voiding during the urodynamic recordings under anesthesia after a large threshold pressure had been achieved. In addition to cystometric deficits in the injured rats, 3 animals displayed uncoordinated EUS EMG activity during the voiding phase where a large increase in tonic EUS EMG activity was observed as the bladder reached its maximum capacity and distension.

**Figure 3.**
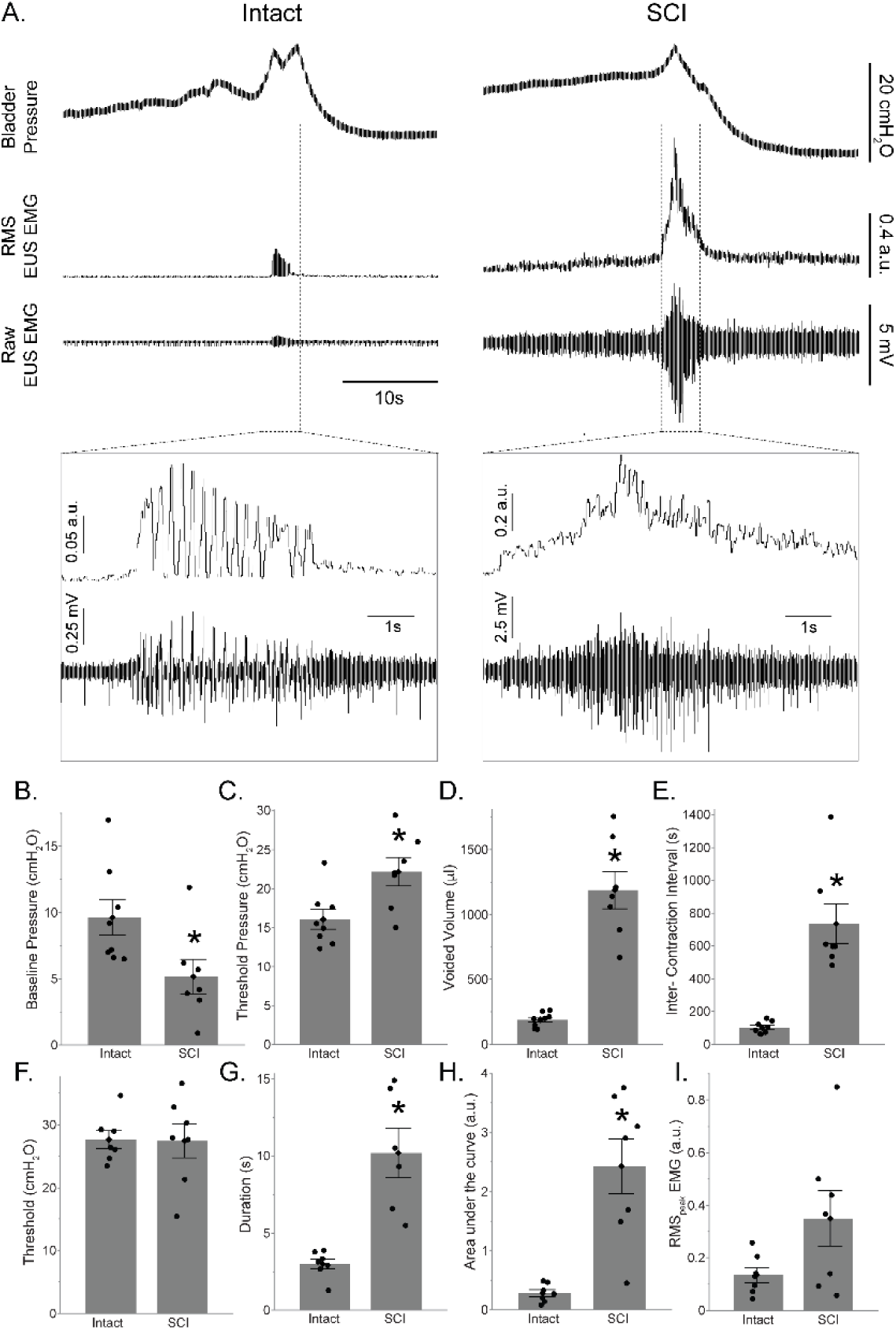
Impact of T9 contusion on cystometric bladder function and EUS EMG activity. A) Example bladder pressure and EUS EMG trace of coordinated voiding in intact animals (left column) and in SCI rats, five days following spinal cord injury (right column). Expanded traces show EUS EMG activity. Note different y-scales in expanded trace used for clarity. (B-E) Summary of the impact of T9 contusion injury on various cystometric outcomes. Baseline pressure was reduced in the SCI group as compared to intact animals. Injured rats had significantly higher threshold pressures, voided volume, and intercontraction intervals. B) Baseline pressure (cmH_2_O); C) Threshold pressure (cmH_2_O); D) Voided volume (μl); E) Intercontraction interval (s). F-I) Summary of the impact of T9 contusion injury on EUS EMG activity. Injured rats had a significantly higher duration, area under the curve, and RMS_peak_ EMG. F) Threshold (cmH_2_O); G) Duration (s); H) area under the curve (arbitrary unit, a.u.); I) RMS_peak_ EMG (a.u.). *p < 0.05. Data are presented as bar plots with all individual data points corresponding to individual animal means. Group means are shown in diamond with error bars depicting ± SE.

Summary urodynamic data highlighting the impact of a T9 contusion on micturition are presented in **Figure 3B-I**. Baseline pressures were significantly lower in the SCI group as compared to the spinal intact group (p = 0.033; **Figure 3B**). In contrast, SCI animals had higher threshold pressures before voiding was initiated as compared to the spinal intact group (p = 0.015; **Figure 3C**). SCI animals also had larger voided volumes (p < 0.001; **Figure 3D**) and exhibited longer periods between contractions (intercontraction interval; p < 0.001; **Figure 3E**). The bladder peak contraction pressures were comparable between spinal intact (29.6 ± 4.5 cmH_2_O, n = 8) and SCI group (30.3 ± 8.1 cmH_2_O, n = 7) (p = 0.83).

There was no difference in the threshold pressure for EUS EMG activation between SCI and spinal intact groups (p = 0.95; **Figure 3F**). The duration of EUS EMG activity was significantly longer in injured rats (p < 0.001; **Figure 3G**). EUS EMG activity in the SCI animals was also characterized by a significant increase in the magnitude of area under the curve (p < 0.001; **Figure H**) and RMS_peak_ EMG tended to be higher in the SCI animals compared to the intact animals (p = 0.0754; **Figure 3I**).

### Effect of Ampakines on Bladder Function

Representative urodynamic recordings following HPCD and ampakine treatment are presented in **Figure 4A**. Mean data for various urodynamic outcomes and treatments are summarized in **Supplemental Table 1**. Parameters of intercontraction interval (**Figure 4B**, F_(4,52)_ = 8.82; *p* < 0.001), voided volume (**Figure 4D**, F_(4,52)_ = 11.71; *p* < 0.001), and threshold pressure (**Figure 4E**, F_(4,52)_ = 19.43; *p* < 0.001), showed a significant group x treatment interaction. Peak pressure showed a treatment effect (**Figure 4C**, *p* < 0.001). *Post-hoc* statistical interactions for group x treatment comparisons are summarized in **Table 1**. Levels not connected with the same letter are significantly different.

**Figure 4.**
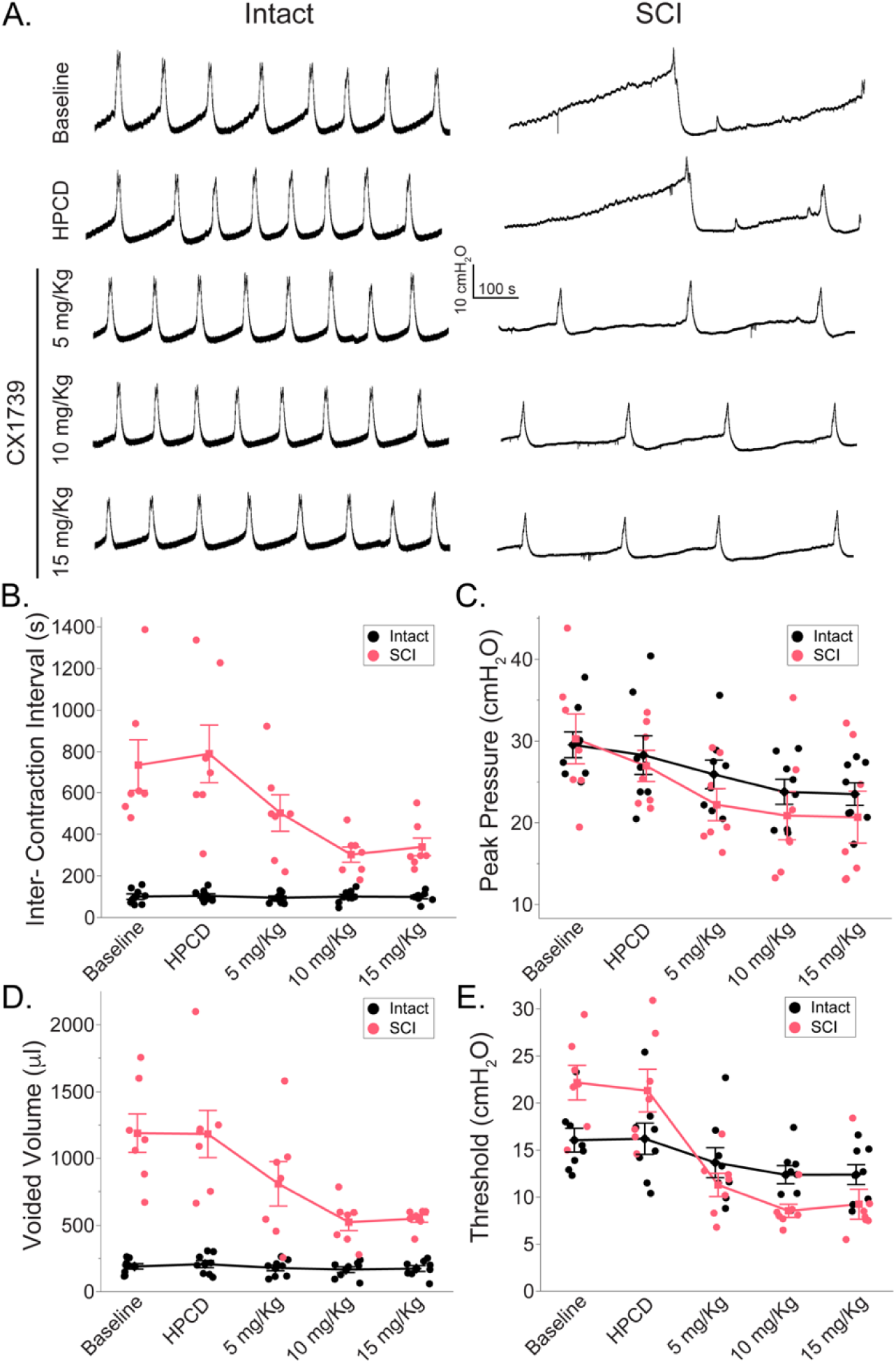
Impact of ampakine treatment on cystometric bladder function. A) Example trace of bladder cystometry in intact animals (left column) and in SCI rats five days following spinal cord injury (right column) following HPCD or ampakine infusion (5 mg/kg, 10 mg/kg, 15 mg/kg). (B-E) Summary of the impact of ampakine treatment on cystometric outcomes in intact (n=8) and SCI (n=7) rats. Ampakine treatment significantly reduced the intercontraction interval, voided volume, and threshold pressure in injured rats. HPCD did not alter cystometry parameters compared to baseline. Ampakine treatment caused a decrease in threshold and peak pressure but did not affect intercontraction interval or voided volume in intact rats. B) Intercontraction interval (s); C) Peak pressure (cmH_2_O); D) Voided volume (μl); E) Threshold pressure (cmH_2_O). Data are presented as line plots with all individual data points corresponding to an individual animal means. Group means are represented with a diamond and error bars depict ± SE.

**Figure 5.**
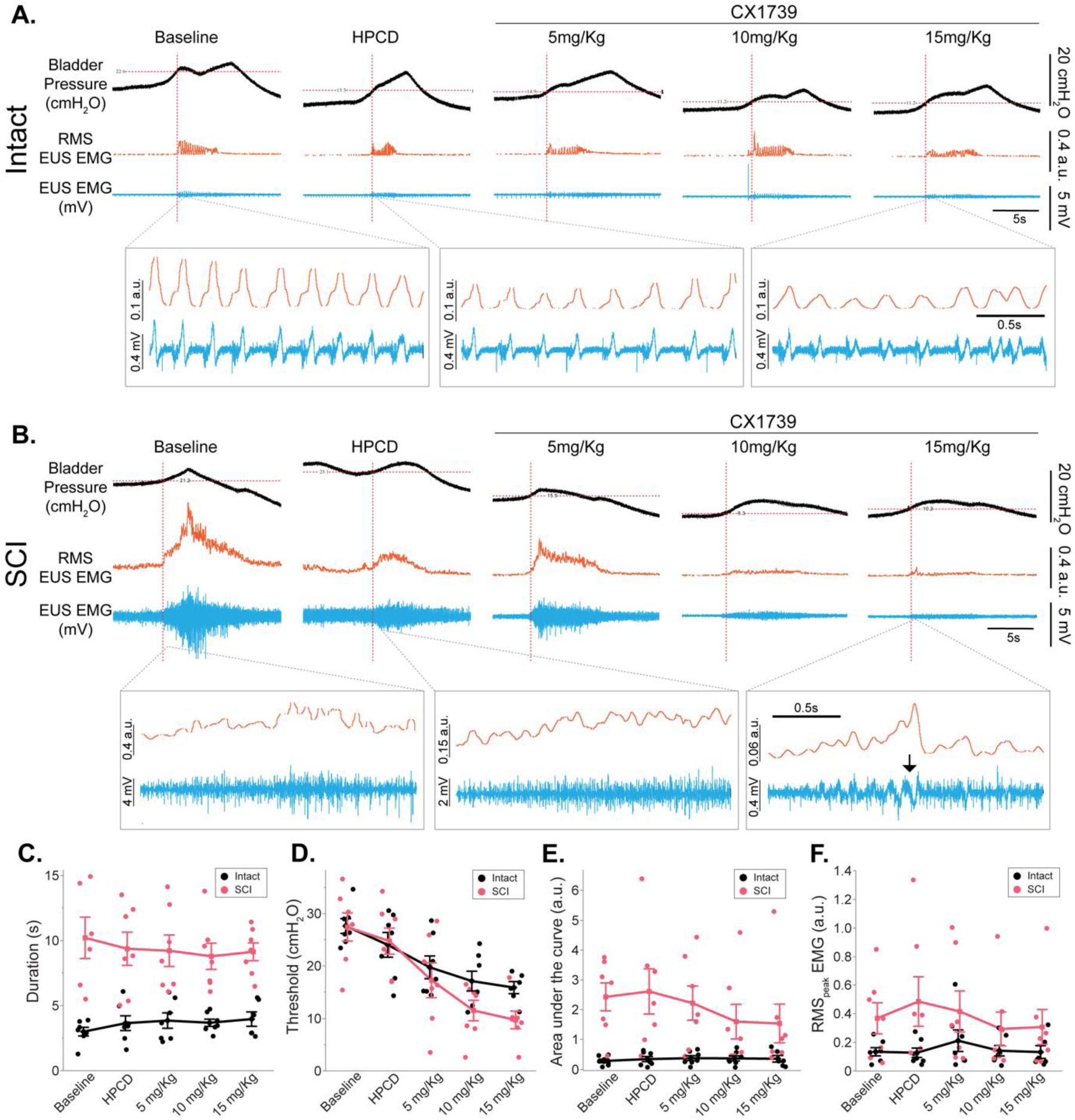
Impact of ampakine treatment on EUS EMG activity. A-B) Example bladder pressure trace, raw EUS EMG trace (blue), and RMS EUS EMG (orange) activity during spontaneous voiding in intact animals and in SCI rats five days following spinal cord injury following HPCD or ampakine infusion (5 mg/kg, 10 mg/kg, 15 mg/kg). Note the re-emergence of coordinated EUS EMG activity following 15 mg/kg ampakine treatment in injured rats (black arrow). (C-F) Summary of the impact of ampakine treatment on EUS EMG activity in intact (n=8) and SCI (n=7) rats. Ampakine treatment reduced the threshold pressure in intact and SCI rats. HPCD had no impact on any outcomes. C) Duration (s); D) Threshold (cmH_2_O; defined as the bladder pressure at which EUS EMG burst was evoked (intersection of red dotted lines)); E) area under the curve (arbitrary unit, a.u.); F) RMS_peak_ EMG (a.u.). Data are presented as line plots with all individual data points corresponding to individual animal means. Group means are represented by a diamond, with error bars depicting ± SE.

**Figure 6.**
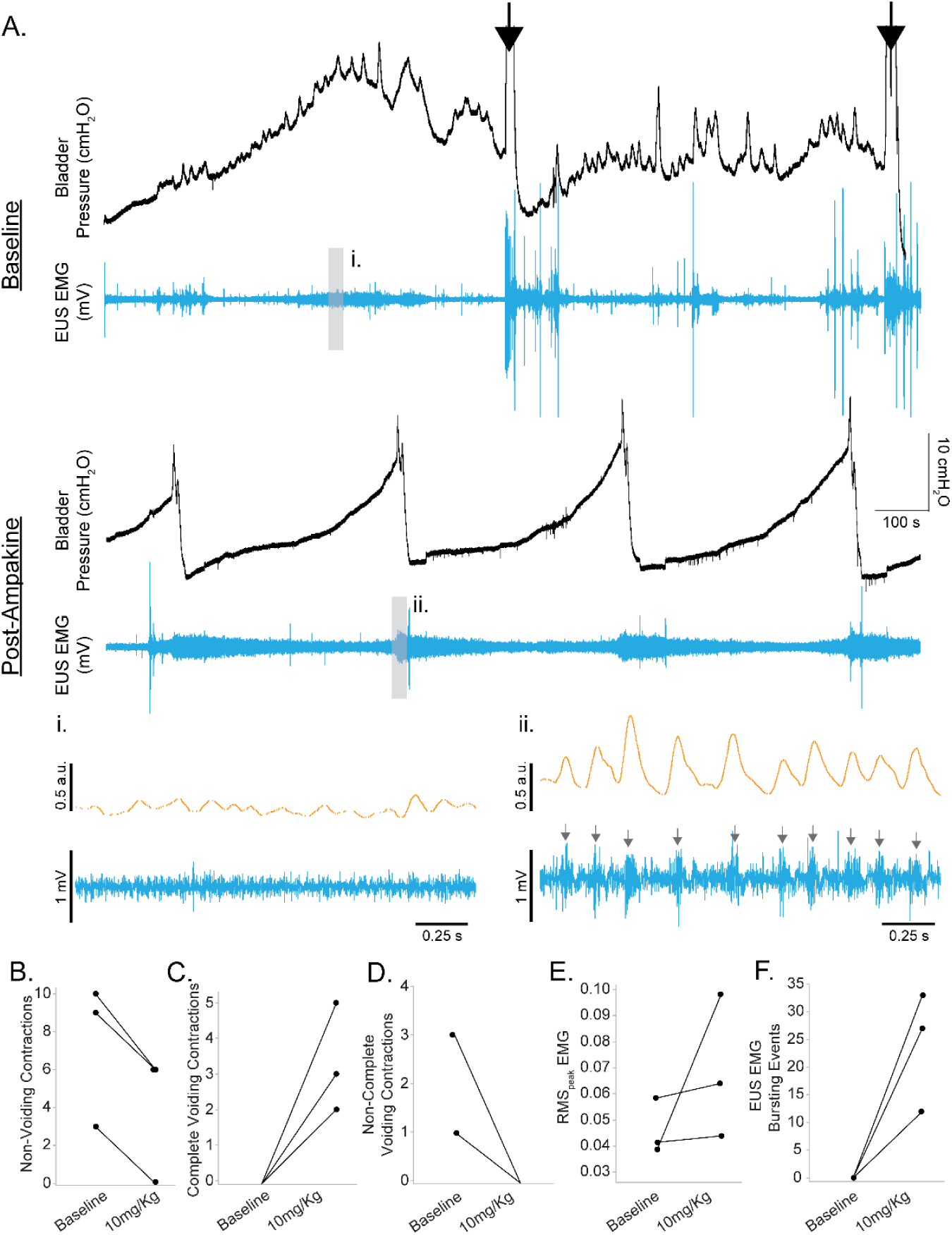
Ampakine rescue coordinated voiding in non-voiding rats following SCI. (**A**) Example bladder pressure and EUS EMG trace of non-coordinated voiding five days following spinal cord injury (Baseline). Black arrows indicate manual expression/emptying of the bladder. The second trace (Post-Ampakine) indicates the same rat following 10 mg/kg CX1739 with voiding re-established. Expanded EUS EMG trace in middle panels shows a lack of coordinated bursting during baseline recording of SCI rats (i). Following ampakine treatment, coordinated EUS bursting is restored (grey arrows; ii). (B-F) Summary of the impact of 10 mg/kg ampakine treatment on cystometry and EUS EMG outcomes in SCI rats displaying non-coordinated voiding (n=3). Ampakine treatment reduced non-voiding and partial voiding contractions in all three rats and established coordinated voiding. No rat exhibited coordinated EUS bursting during the voiding phase at baseline. Ampakine treatment re-established coordinated EUS bursting in all three rats. B. Non-voiding contractions; C) Complete voiding contractions; D) Non-complete voiding contractions; E) RMS_peak_ EMG; F) EUS EMG Bursting Events. Data are presented as individual data points corresponding to individual animal means.

**Table 1.**
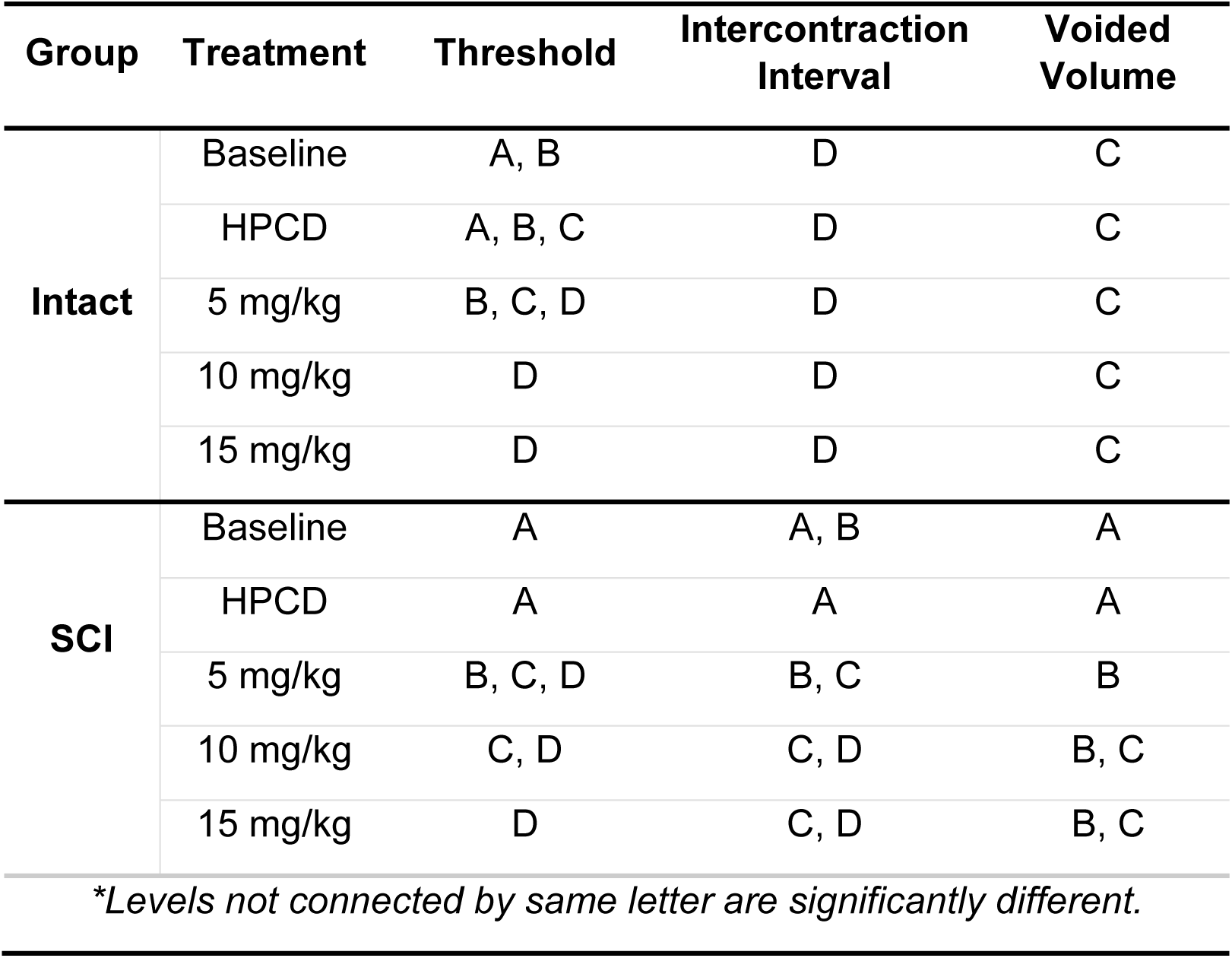
Statistical post-hoc comparisons for cystometric outcomes with significant treatment and group interactions in spontaneously voiding intact (n=8) and SCI (n=7) rats following ampakine treatment.

| Group | Treatment | Threshold | Intercontraction Interval | Voided Volume |
| --- | --- | --- | --- | --- |
| **Intact** | Baseline | A, B | D | C |
|  | HPCD | A, B, C | D | C |
|  | 5 mg/kg | B, C, D | D | C |
|  | 10 mg/kg | D | D | C |
|  | 15 mg/kg | D | D | C |
| **SCI** | Baseline | A | A, B | A |
|  | HPCD | A | A | A |
|  | 5 mg/kg | B, C, D | B, C | B |
|  | 10 mg/kg | C, D | C, D | B, C |
|  | 15 mg/kg | D | C, D | B, C |
| *\*Levels not connected by same letter are significantly different.* |  |  |  |  |

Spinal intact rats demonstrated no effect of HPCD or ampakine treatment on the intercontraction interval, or voided volume. However, 10 and 15 mg/kg ampakine treatment significantly reduced peak pressures in spinal intact rats (28.3 ± 6.7 cmH_2_O with HPCD vs 23.8 ± 4.3 cmH_2_O with 10 mg/kg ampakine and 23.5 ± 3.9 cmH_2_O with 15 mg/kg ampakine, n = 8 (treatment effect; p < 0.001, **Figure 4C**). Ampakine treatment with doses of 10 and 15 mg/kg caused an ∼ 25% reduction in the threshold pressure in these rats (treatment effect, p<0.001, **Figure 4E**).

In rats with SCI, infusion of the HPCD vehicle had no discernable impact on intercontraction interval, peak pressure, voided volume, and threshold pressure. In sharp contrast, the 10 and 15 mg/kg ampakine doses caused an ∼55% decrease in the intercontraction interval (treatment effect; p < 0.001, **Figure 4B**). Similarly, to spinal intact rats, peak pressure decreased in the 10 mg/kg and 15 mg/kg ampakine treated SCI rats (27 ± 5 cmH_2_O with HPCD vs 20.9 ± 7.8 cmH_2_O with 10 mg/kg ampakine and 20.7 ± 8.4 cmH_2_O with 15 mg/kg ampakine, n = 7 (treatment effect; p < 0.001; **Figure 4C**). As compared to the baseline and HPCD treatment, each of the three ampakine doses also resulted in a reduction in voided volume (∼32-58%, treatment effect, p < 0.001, **Figure 4D**.), likely owing to the shorter intercontraction intervals and higher frequency of voiding. Similarly, 5, 10, and 15 mg/kg ampakine doses caused a reduction (∼50-58%, treatment effect, p < 0.001, **Figure 4E**) in threshold pressure as compared to the baseline and HPCD pressure.

### Effects of Ampakine on EUS EMG Activity

Representative EUS EMG traces following HPCD and ampakine treatment are presented in **Figure 5A-B**. Mean data for various EUS EMG parameters at baseline and following treatments are summarized in **Supplemental Table 2**. The EUS burst duration was considerably longer in injured rats as compared to spinal intact rats (**Figure 5C**, *p* < 0.001). Neither HPCD nor ampakine treatment had any effect on EMG burst duration in injured or spinal intact rats (treatment effect; p = 0.52). However, there was a robust treatment effect on the threshold bladder pressure for evoking EUS EMG activity (**Figure 5D**, *p* < 0.001). In both injured and spinal intact rats, ampakine treatment caused EUS EMG bursting to be evoked at much lower bladder pressures as compared to baseline. This effect was also dose-dependent as the 10mg/kg and 15mg/kg doses elicited a larger drop in threshold levels. Ampakine or HPCD treatment had no effect on the magnitude of area under the curve (**Figure 5E**, p = 0.22) or RMS_peak_ EMG (**Figure 5F**, p = 0.20).

### Effects of Ampakine on Non-voiding Rats 5 Days Following SCI

A small cohort of animals (3 of our 10 rats) could not establish a regular cystometric voiding pattern, as evidenced by many non-voiding contractions and slow leaking once the bladder was full (**Figure 6A**). Manual expression of the bladder had to be performed in order to void the bladder during cystometry. This group of animals was assessed separately from the animals displaying spontaneous voiding since parameters of intercontraction interval, base pressure, threshold pressure or peak pressure could not be measured during baseline conditions in the non-voiding animals. Ampakine treatment (10 mg/kg) was successful in reducing the number of non-voiding and partial contractions in all three rats and increasing the frequency of complete voiding contractions (**Figure 6B-D**). Interestingly, we observed the reappearance of EUS EMG bursting events in SCI rats with inefficient coordinated voiding by administration of CX1739 (**Figure 5B**) and clear coordinated bladder contraction with EUS EMG bursting activity with CX1739 in all three SCI rats where there were complete loss of coordination and failed to produce voiding (**Figures 6E** and **6F**). These results suggest that ampakines potentiate micturition circuits in SCI rats at a level that was able to produce a coordinated void.

## Discussion

We demonstrate that systemic (*i.v.*) delivery of a positive allosteric modulator of AMPA receptors, ampakine CX1739, rapidly improves micturition reflexes in sub-acute spinally injured rats. In particular, acute ampakine treatment reduced SCI-induced deficits in the inter-contraction interval, threshold pressure and peak pressure. A decrease in voided volume was also observed with the ampakine treatment, which would reduce the potential for kidney damage in SCI rats. Ampakine treatment reduced the pressure threshold of EUS bursting initiation during the micturition cycle. Ampakines have been safely administered in human clinical trials for other purposes (Boyle et al., 2012; Wesensten et al., 2007), and accordingly, our results suggest that ampakine pharmacotherapy may be a new strategy to support the recovery of bladder function following acute spinal cord injuries.

### Therapeutic Impact of Ampakines

Ampakines are designed to enhance AMPA-mediated glutamatergic neurotransmission (Arai and Kessler, 2007; Lynch, 2006). *In-vitro* studies confirm that ampakines are not AMPA receptor agonists but are allosteric modulators of their activity (Arai et al., 1996; Arai et al., 2002; Arai et al., 2004). Ampakines do not impact NMDA or kainite receptors directly (Lynch, 2006). Importantly, ampakines exist in two distinct classes: high and low-impact ampakines. High-impact ampakines act mainly by changing AMPA receptor desensitization or changing the affinity of an agonist binding to the receptor (Arai et al., 2002). There have been concerns with the use of high-impact ampakines, which can cause hyper-excitability of neuronal circuits and might be limited by a narrow therapeutic window. The mode of action of low-impact ampakines is primarily via changing the rate at which channels open, resulting in a small increase in current amplitude without prolonging channel deactivation (Arai et al., 2002). The positive effects of low-impact ampakines in rodent models (Arai and Kessler, 2007; Lynch, 2006; Lynch and Gall, 2006), without the convulsant effects (Shaffer et al., 2013) of high-impact ampakines make them a better therapeutic candidate. For this reason, our group’s current and prior studies have focused on low-dose, low-impact ampakines following SCI.

We have recently demonstrated that low-impact ampakines can enhance respiratory drive following SCI (Rana et al., 2021; Wollman et al., 2020). High and mid-cervical SCIs result in respiratory compromise due to disrupted glutamatergic pathways that activate motor neurons controlling the diaphragm muscle. AMPA receptors play a large role in mediating this respiratory drive (Thakre et al., 2023). In acute and sub-acute stages following a high cervical SCI, systemic delivery of a low-dose ampakines (CX1739 and CX717, 5 mg/kg) was sufficient to enhance diaphragm EMG activity and overall ventilation in freely behaving rats. Further, systemic delivery of CX1739 or CX717 at 5 mg/kg resulted in no detectable off-target effects, such as changes in heart rate, blood pressure, or overall animal activity. In addition to its efficacy in improving respiratory function in rodent models of SCI, CX1739 was safely administered with no adverse events as part of phase 2 clinical trials to alleviate opioid-induced respiratory depression (RespireRx and University, 2016).

While pharmacokinetic studies were not conducted in the present study, the mean plasma half-life of a single CX1739 intravenous bolus in Sprague Dawley rats is 1.25 ± 0.03 hrs, with a T_max_ (time to maximum plasma) concentration of 30 minutes (information provided through personal communication with RespireRx). The mean range of CX1739 half-life in humans has been reported in the range of 6.6 to 9.3 hours. In our studies, we continued to observe an effect of ampakines for the duration of the 45 minutes post-drug administration. Although the current acute preparations were constrained by time to reduce variability in cystometry recordings and keep the total duration of the preparation under 6 hours, future studies will aim to test the long-term impact of an acute and optimal ampakine dose in awake rats, a critical step along the translational pathway.

Lastly, in the present study, only female rats were studied due to the lower incidence of bladder complications following a thoracic contusion injury. Future studies will need to be completed in both male and female rats to compare potential sex differences in ampakine effects. While we have not observed any sex effects of ampakines in our respiratory studies (Rana et al., 2021) there are known differences in recovery and injury manifestation between male and females that need to be further studied (Anderson et al., 2023; Myers et al., 2023).

### Role of AMPA Receptors in Micturition Circuits

Glutamate signaling through the AMPA-type receptors plays a critical role in several micturition circuits under normal conditions and after SCI (de Groat et al., 2015; Fowler et al., 2008). In general, glutamate is excitatory in bladder circuits, and AMPA receptors play a major role in mediating glutamatergic drive. Indeed, AMPA receptor antagonists can decrease bladder contraction frequency and reduce EUS EMG activity (Yoshiyama et al., 1997). These actions occur via the descending input from the pontine micturition center to the pre-ganglionic detrusor and urethral sphincter motor neurons. AMPA receptors are also expressed in bladder sensory circuits, where inhibition of AMPA receptors reduces sensory input to the brain stem (Kakizaki et al., 1998). They are presumable present on the first-order spinal cord neurons responding to glutamate released by sensory neurons, but this has yet to be proven definitively.

Based on data from the current study, the increased incidence of coordinated bladder voiding in injured rats following systemic ampakine administration would support a centrally mediated pathways involving both the sensory and motor systems (Kakizaki et al., 1998; Matsumoto et al., 1995a, b). The ascending and descending pathways express different AMPA receptor subunits (Shibata et al., 1999). Specifically, dorsal laminae in the L6-S1 rat spinal cord constitute a high abundance of GluR1 and GluR2 subunits (Shibata et al., 1999). Whereas, neurons located in the ventral horn have a high abundance of GluR3 and GluR4 subunits. In pontine micturition centers, high levels of GluR1 and GluR2 subunits have been observed (Shibata et al., 1999). Positive allosteric modulators have been shown to bind at the interface between two adjacent subunits at the D1 lobe of the ligand-binding domain (Jin et al., 2005). Alternative splicing of amino acids at the ligand-binding domain of various receptor subunits can change the domain interface, thereby changing the interaction of allosteric modulators and how they impact channel kinetics and their sensitivity (Golubeva et al., 2022). Based on current data, we presume that ampakines act similarly on various AMPA receptor subunits. However, future studies will determine the location of ampakines action to modulate bladder function following incomplete SCI.

Unlike previous studies using AMPA receptor antagonists, we did not observe any statistically significant effects of CX1739 on bladder contraction or EUS EMG activity in spinally intact rats. Previous studies in naïve rats have shown that AMPA receptor antagonism with LY215490 inhibits detrusor and EUS activity (Yoshiyama et al., 1997). It is important to note that in our study, we used CX1739, which is not an AMPA receptor agonist but an allosteric modulator of the receptor that can enhance endogenous signaling. The AMPA antagonist used in the previous study, LY215490, antagonizes glutamate to bind with postsynaptic AMPA receptors and, thereby, completely inhibiting the glutamatergic synaptic transmission of micturition circuits, resulting in complete inhibition of bladder and EUS activity. In comparison, CX1739 is an allosteric modulator of AMPA receptors and does not directly activate the receptor. Thus, our results indicate that allosterically modulating the receptor with a low impact ampakine does not enhance normal glutamatergic synaptic transmission in spinal intact rats, while it can enhance activity following SCI. Future research is needed to determine why this is the case but it could be due to down-regulation of AMPA receptors (Grossman et al., 1999) and increased release of glutamate in SCI models (Demediuk et al., 1989; Liu et al., 1991; Panter et al., 1990).

### Role of AMPA Receptors in Micturition Circuits after SCI

After SCI, the bladder is initially in an “areflexive state,” and micturition is impaired. As spinal circuits undergo post-SCI neuroplastic changes, the bladder transitions to a neurogenic detrusor overactivity (de Groat et al., 2015; de Groat and Yoshimura, 2006). This results in a hyper-reflexive state and causes sphincter dyssynergia, reflex incontinence, and increased residual urine in the bladder following a void. It is well established that glutamatergic signaling and expression of AMPA receptors are profoundly altered after SCI (Grossman et al., 2001; Grossman et al., 1999; Pikov and Wrathall, 2002; Rana et al., 2022; Yoshiyama et al., 1999a). In the thoracic spinal cord, there is a decrease in the expression of AMPA receptor subunits acutely (e.g., 24 hrs) after thoracic SCI (Grossman et al., 1999). However, AMPA receptor expression increases at more chronic post-SCI time points (e.g., 1 month) following the injury (Grossman et al., 1999). Similar dynamic changes in AMPA receptors have been described in other spinal segments after SCI (Rana et al., 2022). An upregulation of AMPA receptors in the chronic stages of injury may, at least in part, be responsible for the hyperactive bladder phenotype that can occur after chronic SCI (Pikov and Wrathall, 2001, 2002; Yoshiyama et al., 1999b). It is unclear why these receptors are upregulated, although this could occur due to homeostatic upregulation (Turrigiano, 2012) secondary to decreased activity in denervated motoneurons due to the SCI-induced interruption of descending tracts.

Our data indicate that ampakine-mediated modulation of AMPA receptors can rescue the areflexive bladder phenotype five days after SCI. At this acute stage post-contusion injury, glutamatergic neurotransmission is partially impaired due to disrupted descending spinal pathways. Specifically, excitatory glutamatergic pathways originating from the pontine micturition centers would be partially disrupted following a T9 contusion injury. Indeed, histological assessment of the spinal cord tissue (**Figure 2**) shows partial disruption of white matter regions that would contain descending lateral corticospinal tracts from the pontine micturition centers (Fowler et al., 2008). In addition, ascending sensory tracts contributing to the sensation of bladder filling would also be disrupted. Thus, in this state of reduced neural drive, ampakines are likely potentiating residual drive to activate the glutamatergic pathways responsible for the areflexive bladder. These effects of ampakines in the acute phase of injury are particularly exciting. In the chronic phase of injury, it is well documented that detrusor overactivity resulting from hyper-activity glutamatergic circuits tends to develop at the 3-4 weeks post-injury time points (Sartori et al., 2022). While ampakines appear to alleviate bladder dysfunction at this acute timepoint post-injury, it is likely that there is a time-dependent efficacy of ampakine therapy that changes from acute to chronic stages of injury, and these aspects remain to be studied.

Globally more than 500,000 people experience an SCI every year, of which 70-84% of patients showed neurogenic bladder dysfunction (de Groat et al., 1990; Kumar et al., 2018). Current therapeutic options for improving bladder dysfunction in SCI patients are not effective and rely primarily on intermittent catheterizations to prevent bladder overfilling and kidney damage. These approaches may further lead to many bladder and kidney function problems (Xiang et al., 2023). In the present study, we show that CX1739 acutely improves hyporeflexive deficits in bladder function in rats following SCI, as measured by cystometry. Cystometry is an excellent tool to evaluate bladder filling and emptying conditions for drug dose responses, as data can be collected from multiple voids during a relatively short drug activation period. This technique does have its limitations as it is conducted under anesthetized conditions and using non-physiologic filling rates. A critical step in future studies, before translation into the clinical population, will be to evaluate overall bladder voiding in freely-moving animals and how bladder function changes in the acute to chronic stages post-injury in ampakine-treated rats.

## Conclusions

We conclude that intravenous delivery of a low-impact ampakine, CX1739, can improve bladder voiding after thoracic SCI. In rats with impaired bladder voiding due to SCI, CX1739 treatment increased the frequency of coordinated voiding and promoted coordinated EUS EMG activity. Importantly, low-impact ampakines, have been safely administered to humans in clinical trials for other indications (Boyle et al., 2012; Wesensten et al., 2007). Accordingly, we suggest that ampakine pharmacotherapy may represent a viable strategy to improve acute hyporeflexive bladder function in persons with SCI.

## Acknowledgments

The authors gratefully acknowledge Dr. Arnold Lippa and RespireRx for supplying the ampakines used in this work. We also thank the Animal Care Services (ACS) staff of the University of Florida for providing oversight for the care and well-being of all animals used in this study. Figure 1 was created using Biorender.com.

## Author Contributions

SR, FA, DF and AM contributed to the conception and design of the study. SR, FA and RM performed data collection and analysis. SR and FA wrote the first draft of the manuscript. SR assembled and designed the figures. AM and DF supervised and acquired funding for this project, and provided final edits of the manuscript. All authors read and approved the submitted version.

## Data Availability

The data supporting this study’s findings are available from the corresponding author upon reasonable request.

## Animal Studies

All the procedures involving rats and tissue performed in this study were approved by the University of Florida Institutional Animal Care and Use Committee and in strict accordance with the US National Institute of Health (NIH) Guide for the Care and Use of Laboratory Animals.

## Significance

There are limited options for patients with recovery of bladder function following spinal cord injury, with most therapies focusing on treating the symptoms, primarily through catheterization. Here we demonstrate that intravenous delivery of a drug which acts as an allosteric modulator of the AMPA type receptor (an “ampakine”) can rapidly improve bladder function following spinal cord injury. The data suggest that ampakines may be a new therapy for early hyporeflexive bladder states following spinal cord injury.

## Abbreviations

AMPA: α-amino-3-hydroxy-5-methyl-4-isoxazolepropionic acid
SCI: Spinal Cord Injury
T9Ct: Unilateral T9 Contusion Injury
EUS: External Urethral Sphincter
EMG: Electromyography
HPCD: 2-hydroxypropyl-beta-cyclodextrin

## References

Anderson, C.E., Birkhauser, V., Liechti, M.D., Jordan, X., Luca, E., Mohr, S., Pannek, J., Kessler, T.M., Brinkhof, M.W.G., 2023. Sex differences in urological management during spinal cord injury rehabilitation: results from a prospective multicenter longitudinal cohort study. Spinal Cord 61, 43–50.

Andersson, K.E., Soler, R., Fullhase, C., 2011. Rodent models for urodynamic investigation. Neurourol Urodyn 30, 636–646.

Arai, A., Kessler, M., Rogers, G., Lynch, G., 1996. Effects of a memory-enhancing drug on DL-alpha-amino-3-hydroxy-5-methyl-4-isoxazolepropionic acid receptor currents and synaptic transmission in hippocampus. Journal of Pharmacology and Experimental Therapeutics 278, 627–638.

Arai, A.C., Kessler, M., 2007. Pharmacology of ampakine modulators: from AMPA receptors to synapses and behavior. Curr Drug Targets 8, 583–602.

Arai, A.C., Xia, Y.F., Rogers, G., Lynch, G., Kessler, M., 2002. Benzamide-type AMPA receptor modulators form two subfamilies with distinct modes of action. Journal of Pharmacology and Experimental Therapeutics 303, 1075–1085.

Arai, A.C., Xia, Y.F., Suzuki, E., 2004. Modulation of AMPA receptor kinetics differentially influences synaptic plasticity in the hippocampus. Neuroscience 123, 1011–1024.

Bourbeau, D., Bolon, A., Creasey, G., Dai, W., Fertig, B., French, J., Jeji, T., Kaiser, A., Kouznetsov, R., Rabchevsky, A., Santacruz, B.G., Sun, J., Thor, K.B., Wheeler, T., Wierbicky, J., 2020. Needs, priorities, and attitudes of individuals with spinal cord injury toward nerve stimulation devices for bladder and bowel function: a survey. Spinal Cord 58, 1216–1226.

Boyle, J., Stanley, N., James, L.M., Wright, N., Johnsen, S., Arbon, E.L., Dijk, D.J., 2012. Acute sleep deprivation: the effects of the AMPAKINE compound CX717 on human cognitive performance, alertness and recovery sleep. Journal of psychopharmacology 26, 1047–1057.

Chitravanshi, V.C., Sapru, H.N., 1996. NMDA as well as non-NMDA receptors mediate the neurotransmission of inspiratory drive to phrenic motoneurons in the adult rat. Brain Res 715, 104–112.

de Groat, W.C., Griffiths, D., Yoshimura, N., 2015. Neural control of the lower urinary tract. Compr Physiol 5, 327–396.

de Groat, W.C., Kawatani, M., Hisamitsu, T., Cheng, C.L., Ma, C.P., Thor, K., Steers, W., Roppolo, J.R., 1990. Mechanisms underlying the recovery of urinary bladder function following spinal cord injury. J Auton Nerv Syst 30 Suppl, S71–77.

de Groat, W.C., Yoshimura, N., 2006. Mechanisms underlying the recovery of lower urinary tract function following spinal cord injury. Prog Brain Res 152, 59–84.

Demediuk, P., Daly, M.P., Faden, A.I., 1989. Effect of impact trauma on neurotransmitter and nonneurotransmitter amino acids in rat spinal cord. J Neurochem 52, 1529–1536.

Fowler, C.J., Griffiths, D., de Groat, W.C., 2008. The neural control of micturition. Nat Rev Neurosci 9, 453–466.

French, J.S., Anderson-Erisman, K.D., Sutter, M., 2010. What do spinal cord injury consumers want? A review of spinal cord injury consumer priorities and neuroprosthesis from the 2008 neural interfaces conference. Neuromodulation 13, 229–231.

Fuller, D.D., Rana, S., Smuder, A.J., Dale, E.A., 2022. The phrenic neuromuscular system. Handb Clin Neurol 188, 393–408.

Golubeva, E.A., Lavrov, M.I., Radchenko, E.V., Palyulin, V.A., 2022. Diversity of AMPA Receptor Ligands: Chemotypes, Binding Modes, Mechanisms of Action, and Therapeutic Effects. Biomolecules 13.

Grossman, S.D., Rosenberg, L.J., Wrathall, J.R., 2001. Relationship of altered glutamate receptor subunit mRNA expression to acute cell loss after spinal cord contusion. Exp Neurol 168, 283–289.

Grossman, S.D., Wolfe, B.B., Yasuda, R.P., Wrathall, J.R., 1999. Alterations in AMPA receptor subunit expression after experimental spinal cord contusion injury. J Neurosci 19, 5711–5720.

Hamid, R., Averbeck, M.A., Chiang, H., Garcia, A., Al Mousa, R.T., Oh, S.J., Patel, A., Plata, M., Del Popolo, G., 2018. Epidemiology and pathophysiology of neurogenic bladder after spinal cord injury. World J Urol 36, 1517–1527.

Jin, R., Clark, S., Weeks, A.M., Dudman, J.T., Gouaux, E., Partin, K.M., 2005. Mechanism of positive allosteric modulators acting on AMPA receptors. J Neurosci 25, 9027–9036.

Kakizaki, H., Yoshiyama, M., Roppolo, J.R., Booth, A.M., De Groat, W.C., 1998. Role of spinal glutamatergic transmission in the ascending limb of the micturition reflex pathway in the rat. Journal of Pharmacology and Experimental Therapeutics 285, 22–27.

Kumar, R., Lim, J., Mekary, R.A., Rattani, A., Dewan, M.C., Sharif, S.Y., Osorio-Fonseca, E., Park, K.B., 2018. Traumatic Spinal Injury: Global Epidemiology and Worldwide Volume. World Neurosurg. 113, e345–e363.

Lin, Y.T., Hsieh, T.H., Chen, S.C., Lai, C.H., Kuo, T.S., Chen, C.P., Lin, C.W., Young, S.T., Peng, C.W., 2016. Effects of pudendal neuromodulation on bladder function in chronic spinal cord-injured rats. J. Formos. Med. Assoc. 115, 703–713.

Liu, D., Thangnipon, W., McAdoo, D.J., 1991. Excitatory amino acids rise to toxic levels upon impact injury to the rat spinal cord. Brain Res 547, 344–348.

Lynch, G., 2006. Glutamate-based therapeutic approaches: ampakines. Curr Opin Pharmacol 6, 82–88.

Lynch, G., Gall, C.M., 2006. Ampakines and the threefold path to cognitive enhancement. Trends Neurosci 29, 554–562.

Matsumoto, G., Hisamitsu, T., de Groat, W.C., 1995a. Non-NMDA glutamatergic excitatory transmission in the descending limb of the spinobulbospinal micturition reflex pathway of the rat. Brain Res 693, 246–250.

Matsumoto, G., Hisamitsu, T., de Groat, W.C., 1995b. Role of glutamate and NMDA receptors in the descending limb of the spinobulbospinal micturition reflex pathway of the rat. Neurosci Lett 183, 58–61.

Mitsui, T., Murray, M., Nonomura, K., 2014. Lower urinary tract function in spinal cord-injured rats: midthoracic contusion versus transection. Spinal Cord 52, 658–661.

Mitsui, T., Neuhuber, B., Fischer, I., 2011. Acute administration of AMPA/Kainate blocker combined with delayed transplantation of neural precursors improves lower urinary tract function in spinal injured rats. Brain Res 1418, 23–31.

Myers, J.B., Stoffel, J.T., Elliott, S.P., Welk, B., Herrick, J.S., Lenherr, S.M., 2023. Sex Differences in Bladder Management, Symptoms, and Satisfaction After Spinal Cord Injury. J. Urol. 210, 659–669.

Panter, S.S., Yum, S.W., Faden, A.I., 1990. Alteration in extracellular amino acids after traumatic spinal cord injury. Annals of Neurology 27, 96–99.

Piatt, J.A., Nagata, S., Zahl, M., Li, J., Rosenbluth, J.P., 2016. Problematic secondary health conditions among adults with spinal cord injury and its impact on social participation and daily life. J Spinal Cord Med 39, 693–698.

Pikov, V., Wrathall, J.R., 2001. Coordination of the bladder detrusor and the external urethral sphincter in a rat model of spinal cord injury: effect of injury severity. J Neurosci 21, 559–569.

Pikov, V., Wrathall, J.R., 2002. Altered glutamate receptor function during recovery of bladder detrusor-external urethral sphincter coordination in a rat model of spinal cord injury. Journal of Pharmacology and Experimental Therapeutics 300, 421–427.

Rana, S., Sieck, G.C., Mantilla, C.B., 2019. Heterogeneous glutamatergic receptor mRNA expression across phrenic motor neurons in rats. J Neurochem.

Rana, S., Sunshine, M.D., Greer, J.J., Fuller, D.D., 2021. Ampakines stimulate diaphragm activity after spinal cord injury. J Neurotrauma.

Rana, S., Zhan, W.Z., Sieck, G.C., Mantilla, C.B., 2022. Cervical spinal hemisection alters phrenic motor neuron glutamatergic mRNA receptor expression. Exp Neurol 353, 114030.

RespireRx, University, D., 2016. Antagonism of Opioid-Induced Respiratory Depression by CX1739 With Preservation of Opioid Analgesia. https://ClinicalTrials.gov/show/NCT02735629.

Sartori, A.M., Hofer, A.-S., Scheuber, M.I., Rust, R., Kessler, T.M., Schwab, M.E., 2022. Slow development of bladder malfunction parallels spinal cord fiber sprouting and interneurons’ loss after spinal cord transection. Experimental Neurology 348, 113937.

Shaffer, C.L., Hurst, R.S., Scialis, R.J., Osgood, S.M., Bryce, D.K., Hoffmann, W.E., Lazzaro, J.T., Hanks, A.N., Lotarski, S., Weber, M.L., Liu, J., Menniti, F.S., Schmidt, C.J., Hajós, M., 2013. Positive allosteric modulation of AMPA receptors from efficacy to toxicity: the interspecies exposure-response continuum of the novel potentiator PF-4778574. Journal of Pharmacology and Experimental Therapeutics 347, 212–224.

Shibata, T., Watanabe, M., Ichikawa, R., Inoue, Y., Koyanagi, T., 1999. Different expressions of alpha-amino-3-hydroxy-5-methyl-4-isoxazole propionic acid and N-methyl-D-aspartate receptor subunit mRNAs between visceromotor and somatomotor neurons of the rat lumbosacral spinal cord. J Comp Neurol 404, 172–182.

Taweel, W.A., Seyam, R., 2015. Neurogenic bladder in spinal cord injury patients. Res Rep Urol 7, 85–99.

Thakre, P.P., Rana, S., Benevides, E.S., Fuller, D.D., 2023. Targeting drug or gene delivery to the phrenic motoneuron pool. J Neurophysiol 129, 144–158.

Thakre, P.P., Sunshine, M.D., Fuller, D.D., 2022. Spinally delivered ampakine CX717 increases phrenic motor output in adult rats. Respir Physiol Neurobiol 296, 103814.

Turrigiano, G., 2012. Homeostatic synaptic plasticity: local and global mechanisms for stabilizing neuronal function. Cold Spring Harb Perspect Biol 4, a005736.

Wesensten, N.J., Reichardt, R.M., Balkin, T.J., 2007. Ampakine (CX717) effects on performance and alertness during simulated night shift work. Aviat Space Environ Med 78, 937–943.

Wollman, L.B., Streeter, K.A., Fusco, A.F., Gonzalez-Rothi, E.J., Sandhu, M.S., Greer, J.J., Fuller, D.D., 2020. Ampakines stimulate phrenic motor output after cervical spinal cord injury. Exp Neurol 334, 113465.

Xiang, L., Li, H., Xie, Q.Q., Siau, C.S., Xie, Z., Zhu, M.T., Zhou, B., Li, Z.P., Wang, S.B., 2023. Rehabilitation care of patients with neurogenic bladder after spinal cord injury: A literature review. World J Clin Cases 11, 57–64.

Yoshiyama, M., 2009. Glutamatergic Mechanisms Controlling Lower Urinary Tract Function. LUTS: Lower Urinary Tract Symptoms 1, S101–S104.

Yoshiyama, M., de Groat, W.C., 2005. Supraspinal and spinal alpha-amino-3-hydroxy-5-methylisoxazole-4-propionic acid and N-methyl-D-aspartate glutamatergic control of the micturition reflex in the urethane-anesthetized rat. Neuroscience 132, 1017–1026.

Yoshiyama, M., Nezu, F.M., Yokoyama, O., Chancellor, M.B., de Groat, W.C., 1999a. Influence of glutamate receptor antagonists on micturition in rats with spinal cord injury. Exp Neurol 159, 250–257.

Yoshiyama, M., Nezu, F.M., Yokoyama, O., Chancellor, M.B., de Groat, W.C., 1999b. Influence of Glutamate Receptor Antagonists on Micturition in Rats with Spinal Cord Injury. Experimental Neurology 159, 250–257.

Yoshiyama, M., Roppolo, J.R., de Groat, W.C., 1997. Effects of LY215490, a competitive alpha-amino-3-hydroxy-5-methylisoxazole-4-propionic acid (AMPA) receptor antagonist, on the micturition reflex in the rat. Journal of Pharmacology and Experimental Therapeutics 280, 894–904.

