## Supplemental Tables for "Acute ampakines increase voiding function and coordination in a rat model of SCI"

***Supplemental Table 1.*** Mean data of cystometry measures at 5- days post-injury following HPCD or ampakine CX1739 treatment. Intact (n = 8), SCI (n = 7) groups. Data are presented as Mean ± SD.

| **Group** | **Treatment** | **Threshold ($cmH_2O$)** | **Intercontraction Interval *(s)*** | **Voided Volume *(μl)*** | **Peak Pressure ($cmH_2O$)** |
|---|---|---|---|---|---|
| **Intact** | Baseline | 16.1 ± 3.6 | 101.7 ± 37 | 189 ± 56.4 | 29.6 ± 4.5 |
| | HPCD | 16.2 ± 4.7 | 105.3 ± 27.9 | 206.8 ± 77.6 | 28.3 ± 6.7 |
| | 5 mg/Kg | 13.7 ± 4.5 | 95.8 ± 25.8 | 178 ± 62.3 | 25.9 ± 5 |
| | 10 mg/Kg | 12.4 ± 2.7 | 101.6 ± 31.6 | 165.1 ± 62.2 | 23.8 ± 4.3 |
| | 15 mg/Kg | 12.4 ± 3 | 99.7 ± 24.1 | 173 ± 61.9 | 23.5 ± 3.9 |
| **SCI** | Baseline | 22.2 ± 4.9 | 734.7 ± 322.1 | 1188.3 ± 381.6 | 30.3 ± 8.1 |
| | HPCD | 21.3 ± 6 | 789 ± 367.3 | 1182.1 ± 467.6 | 27 ± 5 |
| | 5 mg/Kg | 11.3 ± 3.3 | 504.2 ± 231.6 | 808.9 ± 440.8 | 22.2 ± 5.2 |
| | 10 mg/Kg | 8.5 ± 1.8 | 303.8 ± 97.4 | 521.3 ± 167.4 | 20.9 ± 7.8 |
| | 15 mg/Kg | 9.2 ± 4.2 | 340.7 ± 113.1 | 548.6 ± 73.8 | 20.7 ± 8.4 |

***Supplemental Table 2.*** Mean data of EUS EMG activity at 5- days post-injury following HPCD or ampakine CX1739 treatment. Intact (n = 8), SCI (n = 7) groups. Data are presented as Mean ± SD.

| Group | Treatment | Duration (s) | Threshold ($cmH_2O$) | Area under the curve *(a.u.)* | $RMS_{peak}$ EMG *(a.u.)* |
|---|---|---|---|---|---|
| **Intact** | Baseline | 3.0 ± 0.9 | 27.6 ± 3.7 | 0.3 ± 0.2 | 0.1 ± 0.1 |
| | HPCD | 3.6 ± 1.5 | 24 ± 6.2 | 0.3 ± 0.2 | 0.1 ± 0.1 |
| | 5 mg/Kg | 3.8 ± 1.6 | 19.7 ± 5.7 | 0.4 ± 0.2 | 0.2 ± 0.2 |
| | 10 mg/Kg | 3.7 ± 0.7 | 17.1 ± 5 | 0.4 ± 0.2 | 0.1 ± 0.1 |
| | 15 mg/Kg | 4.2 ± 1.3 | 15.6 ± 2.7 | 0.4 ± 0.3 | 0.1 ± 0.1 |
| **SCI** | Baseline | 10.2 ± 3.9 | 27.4 ± 7.1 | 2.4 ± 1.2 | 0.3 ± 0.3 |
| | HPCD | 9.4 ± 3.4 | 24.7 ± 6.6 | 2.6 ± 2 | 0.5 ± 0.4 |
| | 5 mg/Kg | 9.2 ± 3.2 | 17.3 ± 8.8 | 2.2 ± 1.5 | 0.4 ± 0.4 |
| | 10 mg/Kg | 8.8 ± 2.7 | 11.5 ± 5.2 | 1.6 ± 1.5 | 0.3 ± 0.3 |
| | 15 mg/Kg | 9.1 ± 1.8 | 9.7 ± 4.4 | 1.5 ± 1.7 | 0.3 ± 0.3 |
